# Hedgehog-Hippo pathway interactions promote T cell exclusion from the tumor microenvironment in basal cell carcinoma

**DOI:** 10.1101/2024.07.07.602398

**Authors:** G. Stockmaier, S. R. Varkhande, P. W. Krenn, D. Hieu-Hoa, A. Sharma, D. P. Elmer, M. Steiner, N. Zaborsky, D. Neureiter, R. Greil, J. Horejs-Hoeck, N. Fortelny, I. K. Gratz, F. Aberger

## Abstract

Basal cell carcinoma (BCC) is the most common non-melanoma skin cancer driven primarily by genetic activation of the Hedgehog/GLI (HH/GLI) signaling pathway. BCC also displays a high mutational burden, with frequent co-occurrence of mutations in the TP53, NOTCH, N-MYC, and Hippo/YAP pathways, though the impact of these mutations on HH/GLI-driven tumorigenesis remains poorly understood. This study examines the interaction between the HH/GLI and Hippo/YAP signaling pathways in the pathogenesis of BCC and the establishment of an immunosuppressive tumor microenvironment (TME). The study used 3D human organotypic skin models and humanized mouse models to demonstrate that co-activation of HH/GLI and Hippo/YAP in epidermal cells suppresses T cell chemotaxis. Transcriptome analysis revealed a significant downregulation of chemotactic and inflammatory genes, particularly under combined GLI-YAP activation. Analysis of primary human BCC biopsies confirmed these findings and demonstrated a marked suppression of T cell chemo-attractants, including CCL22 and CCL27. Spatial profiling of immune cells in BCC tissues revealed preferential exclusion of CD8^+^ T cells from the TME, which correlated with high HH/GLI and Hippo/YAP activity. Further, functional assays in humanized mice showed that the GLI-YAP interaction controls immune cell distribution by inhibiting T cell recruitment and migration into the skin. These findings suggest that cooperation between oncogenic HH/GLI and Hippo/YAP signaling contributes to the development of an immunosuppressive TME in BCC by disrupting local chemokine signaling. Understanding these mechanisms provides a basis for the development of targeted combination therapies to enhance the efficacy of existing treatments, such as Smoothened (SMO) inhibitors and immune checkpoint blockers, by promoting T cell infiltration and activation within the TME.

## Introduction

Basal cell carcinoma (BCC) is the most common non-melanoma skin cancer with a high mutational burden. In the U.S. alone, approximately 3-4 million new cases of BCC are diagnosed each year^1–3^. Genetic lesions, such as those caused by chronic UV irradiation over an individuaĺs lifetime, account for a spectrum of recurrent driver mutations not only in the main tumor promoting Hedgehog/GLI (HH/GLI) pathway, but also in TP53, Hippo/YAP, N-MYC and/or NOTCH signaling^1^. This complex mutational landscape drives malignant disease progression including the establishment of an immunosuppressive tumor microenvironment (TME). However, our understanding of the contributions of individual genetic alterations to HH/GLI-induced BCC pathogenesis in relation to immune regulative processes is largely incomplete^1, 4^.

Mutational activation of HH/GLI signaling is the primary genetic driver event implicated in more than 90% of BCC patients. Most BCC display aberrant HH/GLI signaling due to loss-of-function mutations in the HH receptor and pathway repressor Patched (PTCH) or activating mutations in the essential HH effector Smoothened (SMO)^1, 5–8^. These mutations lead to constitutive activation of Glioma-Associated Oncogene (GLI) transcription factors at the distal end of the HH pathway, resulting in BCC development and growth^9^. In addition, the oncogenicity of HH/GLI is further modulated and enhanced by crosstalk with other oncogenic signals and driver mutations^10–17^. Among these, Hippo pathway components have been shown to be mutated in approximately 50% of BCC patients, resulting in enhanced activity of the downstream transcription factor Yes-Associated Protein 1 (YAP1)^1, 10, 18, 19^. Notably, YAP activity has been shown to be essential for HH/GLI-driven BCC initiation and progression in mouse models, but little is known about the mechanistic role of activated Hippo/YAP signaling and its interaction with HH/GLI in regulating a tumor-promoting microenvironment and evading the anti-tumor immune response, two processes of high therapeutic relevance^19, 20^.

The causal role of HH/GLI signaling in BCC development led to the identification and FDA approval of vismodegib and sonidegib, two small-molecule inhibitors of the essential HH effector SMO, for the treatment of metastatic (mBCC) and locally advanced (laBCC) BCC^21–24^. Therapeutic responses are seen in approximately 50% of sporadic BCC patients, but are not durable as patients develop drug resistance, or discontinue treatment due to severe adverse events^25, 26^. More recently, several anti-PD1 immune checkpoint blockers (ICBs) including pembrolizumab, nivolumab and cemiplimab - some in combination with vismodegib - have been evaluated in clinical trials^26–29^. This has led to the approval of cemiplimab for the treatment of mBCC or laBCC resistant or unresponsive to SMO inhibitor treatment. However, the overall response rate to ICB therapy in BCC patients ranged from only 22% to 38%, calling for the improvement of treatment strategies^26–29^. Therefore, a better understanding of the molecular processes and signals that control the formation of an immunosuppressive TME, including the cues and mechanisms that regulate immune cell infiltration and their intra-tumoral spatial distribution, is crucial for the development of more effective combination therapies.

In this study, we investigated the functional consequences of the joint activation of HH/GLI and Hippo/YAP signaling in the context of BCC formation and its impact on the tumor immune microenvironment. Using human organotypic *in vitro* and humanized murine xenografting *in vivo* models, we show that combined activation of GLI-YAP signaling in epidermal cells establishes an immunosuppressive microenvironment that limits T cell infiltration by modulating the cytokine and chemokine milieu in the TME. These data may provide the basis for targeted combination therapies to enhance the effect of approved SMO inhibitors and ICBs.

## Results

### Antagonistic GLI-YAP interactions in epidermal cells repress chemokine signaling and lymphocyte chemotaxis genes

Activating mutations in HH/GLI and Hippo/YAP signaling are highly prevalent in tumors of BCC patients and frequently co-occurring (suppl. Fig. S1A). To mimic and study the consequences of co-activation of oncogenic HH/GLI and Hippo/YAP signaling in BCC, we established 3D human organotypic skin cultures (OTCs) by retrovirally overexpressing either active GLI2 (G^act^)^30^ and active YAP1 mutant YAP^S127A^ (Y^act^) alone or in combination (GY^act^) in human keratinocytes (N/TERT, Fig. 1A, left), which are capable of forming a stratified epidermis in 3D OTCs when co-cultured with human fibroblasts in an air-liquid interface setup for two weeks (Fig. 1A, middle)^31, 32^. Oncogene expression in 2D and the epithelial compartment in 3D OTCs was verified at the protein and mRNA level (suppl. Fig. S1B, C). Notably, we observed histological changes in skin architecture and morphology in response to G^act^ and Y^act^ expression, of which G^act^ overexpression induced hyperproliferative nests, while GY^act^ also caused subtle budding and epithelial downgrowth reminiscent of invasive processes. This suggests distinct biological activity of the genetic alterations introduced into the epidermal compartment of 3D OTCs (Fig. 1B). After epidermal stratification of the 3D OTCs, we separated the epithelial from the dermal compartment via enzymatic digestion and subjected the epidermal samples to RNA-sequencing (RNA-seq) analysis to identify cell-autonomous effects of single and combined activation of HH/GLI and Hippo/YAP signaling (Fig.1A right).

**Figure 1:**
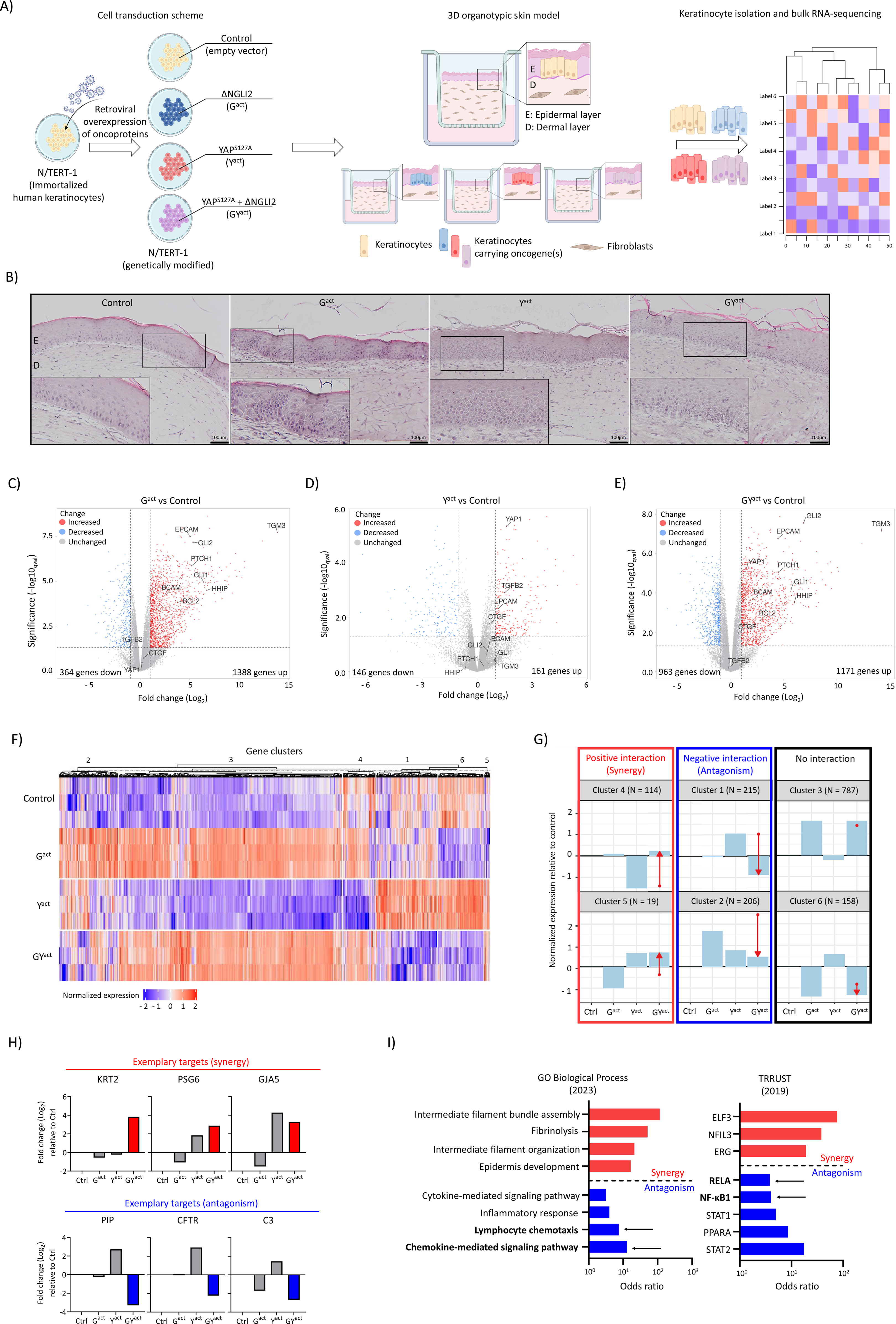
Oncogenic HH-HIPPO signal interactions in 3D organotypic skin cultures. **A)** Illustration of human immortalized keratinocytes (N/TERT-1) and retroviral transduction scheme with empty control vectors (termed control), HA-tagged ΔNGLI2 (G^act^), FLAG-tagged YAP^S127A^ (Y^act^), or the combination of G^act^ and Y^act^ (GY^act^). Dermal matrix consisted of fibrinogen and human immortalized fibroblasts. Keratinocytes overexpressing oncogenes were placed on top to grow 3D human organotypic skin cultures (OTCs) under air-liquid interface conditions for 14 days. The epidermis was enzymatically separated from the dermis for bulk RNA-seq. **B)** Representative Hematoxylin-Eosin (H&E) stainings of control, G^act^, Y^act^, GY^act^ 3D OTCs. E: Epidermis, D: Dermis. Scale bars 100 µm. **C-E)** Volcano plots showing DEGs (Log_2_ fold-change > 1 or < −1, q_val_ < 0.05) for G^act^, Y^act^, and GY^act^ vs control 3D OTC epidermis. Red dots indicate genes upregulated by the specific oncogene(s); blue dots represent downregulated genes. **F)** Heatmap of DEGs (Log_2_ fold-change > 1 or < −1, q_val_ < 0.05, resulting in 1499 DEGs) comparing all four conditions (control, G^act^, Y^act^, GY^act^). Hierarchical clustering revealed six distinct clusters of genes with similar expression behavior, containing in total 1499 DEGs. **G)** Combined gene expression pattern of all genes per cluster from (F). “N” refers to the number of genes per cluster. Length and direction of the red arrow indicate the magnitude and direction of GLI-YAP interaction effects. Left (red frame): clusters and genes under positive interaction (synergy, cluster 4 and 5, 133 genes total); middle (blue frame): negative interaction (antagonism, cluster 1 and 2, 421 genes total); right (black frame): no interaction (Cluster 3 and 6, 945 genes total). Y-axis: normalized expression relative to control. **H)** Gene expression behavior of selected genes under positive (synergy, top) or negative (antagonism, bottom) GLI-YAP control. Y-axis: Fold change (Log_2_) relative to control. **I)** Bar chart of significantly enriched (q_val_ < 0.05) GO BP and TRRUST terms of all genes under synergistic (133 genes) or antagonistic (421 genes) GLI-YAP interaction effects ranked by odds ratio.

Bioinformatic analysis of RNA-seq data from the epidermal compartment revealed the regulation of known HH/GLI and Hippo/YAP target genes. Specifically, G^act^ expression in keratinocytes induced a BCC-like gene signature with an expected upregulation of HH target genes (GLI1, GLI2, PTCH1, HHIP) and of typical BCC biomarkers (EPCAM, BCAM, BCL2, TGM3) (Fig. 1C), resembling the molecular nature of human BCC^1, 2, 15, 71^. In contrast to G^act^, Y^act^ expression in the epidermal compartment of 3D OTCs resulted in the upregulation of YAP target genes such as TGFB2 and CTGF. Y^act^ did not induce the expression of HH targets or BCC biomarkers, except for EPCAM (Fig. 1D). GY^act^ epidermis exhibited both HH/GLI and Hippo/YAP target gene expression, confirming concomitant pathway activity. Interestingly, GY^act^ co-expression resulted in far more genes downregulated than G^act^ or Y^act^ expression alone, suggesting potential interaction effects of the two oncogenes repressing gene expression in transformed keratinocytes (Fig. 1E). To explore the intricacies of GLI- YAP interaction effects at a global level, all 1499 differentially expressed genes (DEGs) between control, G^act^, Y^act^, and GY^act^ were hierarchically clustered to reveal the prevailing gene expression patterns, resulting in six clusters (Fig. 1F). When analyzing for possible interaction (more than additive) effects between GLI-YAP, we found two clusters (4 and 5, with a total of 133 genes) with positive interaction effects (synergy) and two clusters (1 and 2, with a total of 421 genes) with negative interaction effects (antagonism). In contrast, approximately two thirds of all genes (945 genes) showed no interaction and were under dominant control of GLI (Fig. 1G-H). A Gene Ontology - Biological Processes (GO BP) and TRRUST enrichment analysis for genes under negative interaction effects (blue) revealed that GLI-YAP antagonism led to reduced inflammatory, chemokine and chemotaxis signaling, potentially via inhibition of RELA/NF-κB1 (Fig. 1I). This included repressed genes such as C3, CCL22, or CCL27, which we confirmed by q-PCR in a larger sample cohort (suppl. Fig. S1D). These data suggest a potential role for GLI-YAP interaction in dampening chemokine signaling, which may contribute to the establishment of an immunosuppressive TME in BCC.

### Reduced chemokine expression profiles correlate with Hedgehog-Hippo activity and low T cell numbers in primary human BCC

To address the potential role of HH/GLI and Hippo/YAP in antagonizing chemokine signaling in primary human BCC, we re-analyzed bulk RNA-seq data of BCC and normal skin biopsies generously provided by Bonilla et al. (2016) (Fig. 2A)^1^. Consistent with the 3D OTC expression data, the comparison between tumor and normal skin revealed a striking downregulation of several chemokines and inflammatory molecules in the tumor biopsies This included the chemokines CCL17, CCL19, CCL22, and CCL27, which are known chemo-attractants for skin-tropic T cells, as well as inflammatory markers involved in tissue-immune crosstalk (SAA1, IL33, IL1A, C3). As expected, typical HH and BCC biomarkers (PTCH1, GLI1, GLI2, HHIP, EPCAM, BCAM, N-MYC, CCND2) were significantly upregulated in tumor samples (Fig. 2B)^33–40^. A detailed, patient-specific analysis of the data revealed that preferentially those BCC samples with high HH/GLI activity showed a strong reduction in the expression of chemotactic and inflammatory genes (Fig. 2C).

**Figure 2:**
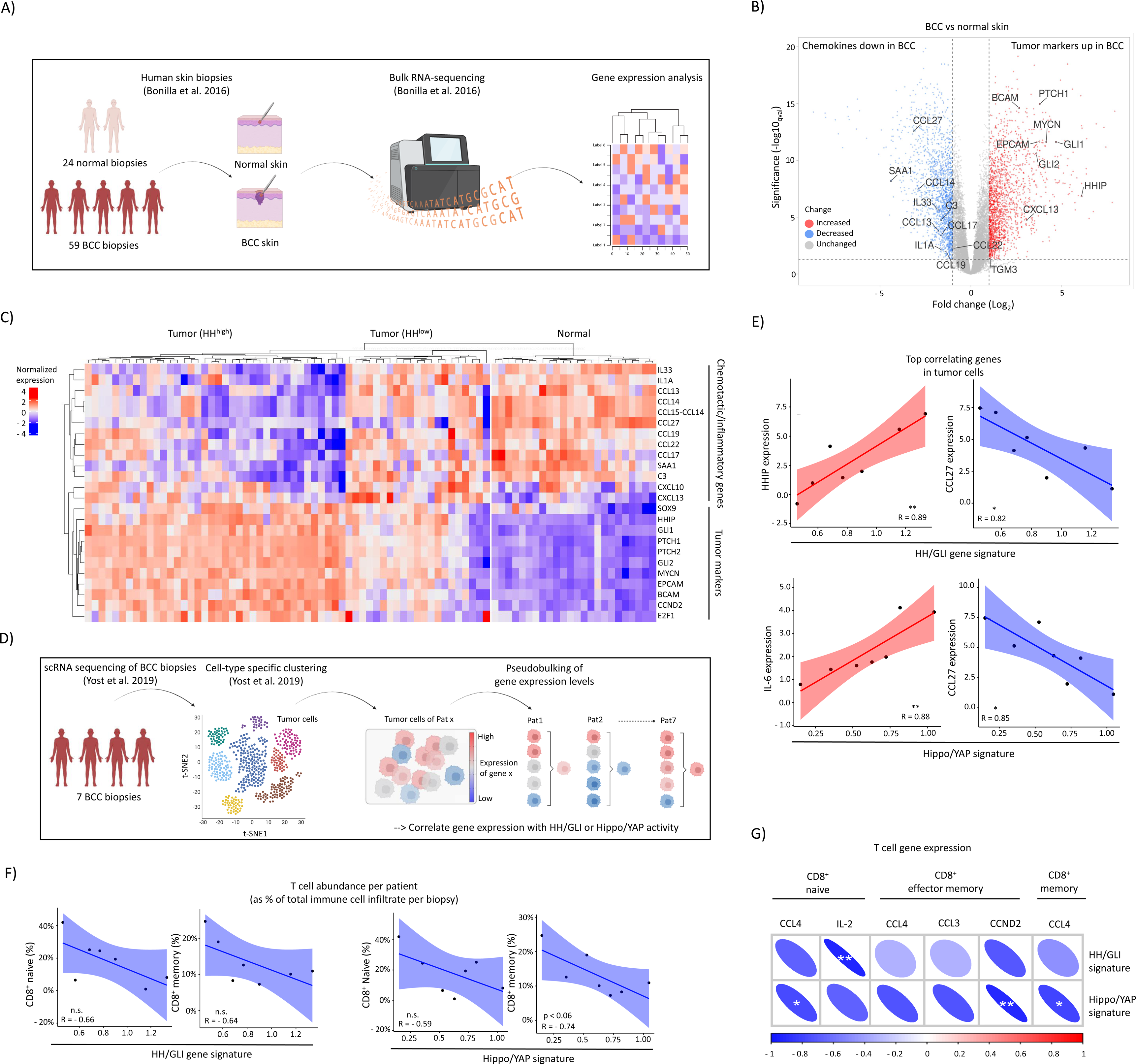
Repression of chemokine expression by GLI-YAP1 interactions in human BCC. **A)** Bulk RNA-seq data from human skin biopsies (24 normal skin and 59 BCC) obtained from Bonilla et al. (2016)^1^ were re-analyzed for DEGs in silico. **B)** Volcano plot showing DEGs (Log_2_ fold-change > 1 or < −1, q_val_ < 0.05) for BCC vs normal skin. Selected upregulated (red) HH and BCC biomarkers and downregulated (blue) chemokines and inflammatory genes are highlighted. **C)** Heatmap of selected genes of interest with dependency on HH/BCC biomarker expression. **D)** Single cell RNA-seq data from human BCC biopsies (N = 7) obtained from Yost et al. (2019)^41^ were re-analyzed for correlation of tumor-derived oncogenic signatures (HH/GLI and Hippo/YAP) with chemokine expression and T cell abundance. **E)** Pearson correlation between HH/GLI (top row) or Hippo/YAP (bottom row) gene signatures and top positive or negative correlating genes in tumor cells of BCC patients. Red: positively correlated, blue: negatively correlated. N = 7. *p<0.05; **p<0.01. **F)** Pearson correlation plots between HH/GLI (left) or Hippo/YAP (right) activity in tumor cells and relative abundance of CD8^+^ naive or CD8^+^ memory T cells compared to other immune cell types in BCC patient biopsies. Blue: negative correlation. N = 7. n.s. (not significant). **G)** Pearson correlation between HH/GLI (top row) or Hippo/YAP (bottom row) activity in tumor cells and gene expression in CD8^+^ T cells. Red: positive correlation; blue: negative correlation. N = 7. *p<0.05; **p<0.01.

To better address a possible link between chemokine downregulation and antagonistic interactions of HH/GLI and Hippo/YAP activity, we re-analyzed single-cell RNA-seq data of BCC biopsies from Yost et al. (2019)^41^. Patients were clustered into high and low expressers based on their HH/GLI (PTCH1, PTCH2, GLI1, GLI2) or Hippo/YAP (TGFB2, CTGF, CYR61, CRIM1, AXL) gene signature in tumor cells, and subsequently screened for positively and negatively correlated genes (Fig. 2D, suppl. Fig. S2A). Consistent with the observed mutational landscape of BCC (suppl. Fig. S1A), we found that HH/GLI and Hippo/YAP activity levels were heterogeneous across patients, though both pathways were positively correlated with each other (R = 0.73, p = 0.062) (suppl. Fig. S2B-C). Looking at the top correlating genes in tumor cells, HH/GLI and Hippo/YAP activity were both strongly positively associated with well-reported target genes (e.g., HHIP and IL-6, respectively). Notably, one of the top negatively linked genes with both pathways was CCL27, which is associated with T cell trafficking towards and within the skin (Fig. 2E). Downregulation of CCL27 in HH- Hippo^high^ BCC tumor cells is in line with its downregulation in bulk RNA-seq data (Fig. 2B) and repression by GLI-YAP co-activation in the epidermis of 3D OTCs (suppl. Fig. S1D). Taken together, these data implicate HH/GLI and Hippo/YAP signaling in the deregulation of the local chemokine milieu in BCC toward a potentially immunosuppressive microenvironment.

To assess whether oncogenic signaling also affects the immune cell composition of patient biopsies, we correlated HH/GLI and Hippo/YAP activity in tumor cells with the relative abundance of immune cells and types within each biopsy. Interestingly, both pathways showed a negative correlation with CD8^+^ memory and, to a lesser extent, also CD8^+^ naive T cell abundance (Fig. 2F). This suggests that patients with higher activity of both pathways had less CD8^+^ T cells in their microenvironment relative to other immune cells, possibly due to reduced chemokine signaling from tumor cells, e.g., reduced CCL27 signaling. When we analyzed the gene expression of these T cell subpopulations in relation to HH/GLI and Hippo/YAP activity in tumor cells, we also found lower levels of the immune cell activation genes CCND2, IL-2, CCL3, and CCL4 produced by these T cells (Fig. 2G).

These findings suggest that the combination of oncogenic HH/GLI and Hippo/YAP signaling in keratinocytes promotes a T cell-hostile TME characterized by inhibition of T cell infiltration and activation due to disruption of chemokine networks in the skin.

### Spatial immune cell profiling of BCC reveals T cell exclusion from the proximal tumor microenvironment

To gain insight into the spatial distribution and activation status of T cells and other immune cell types within the TME of BCC, we performed spatial profiling of immune cell markers in formalin-fixed, paraffin-embedded (FFPE) BCC patient material by digital spatial profiling technology using GeoMx DSP coupled to nCounter systems (Nanostring). Specifically, we analyzed the protein expression of CD3, CD8, CD4, CD45, ß2-Mi, HLA-DR, CD11c, CD68, CD56, GZMB, Ki-67, PD-1, and CTLA-4 in spatially distinct CD45 positive “immune” regions (N = 6) per BCC patient sample (N = 3) (Fig. 3A, suppl. Fig. S3A).

**Figure 3:**
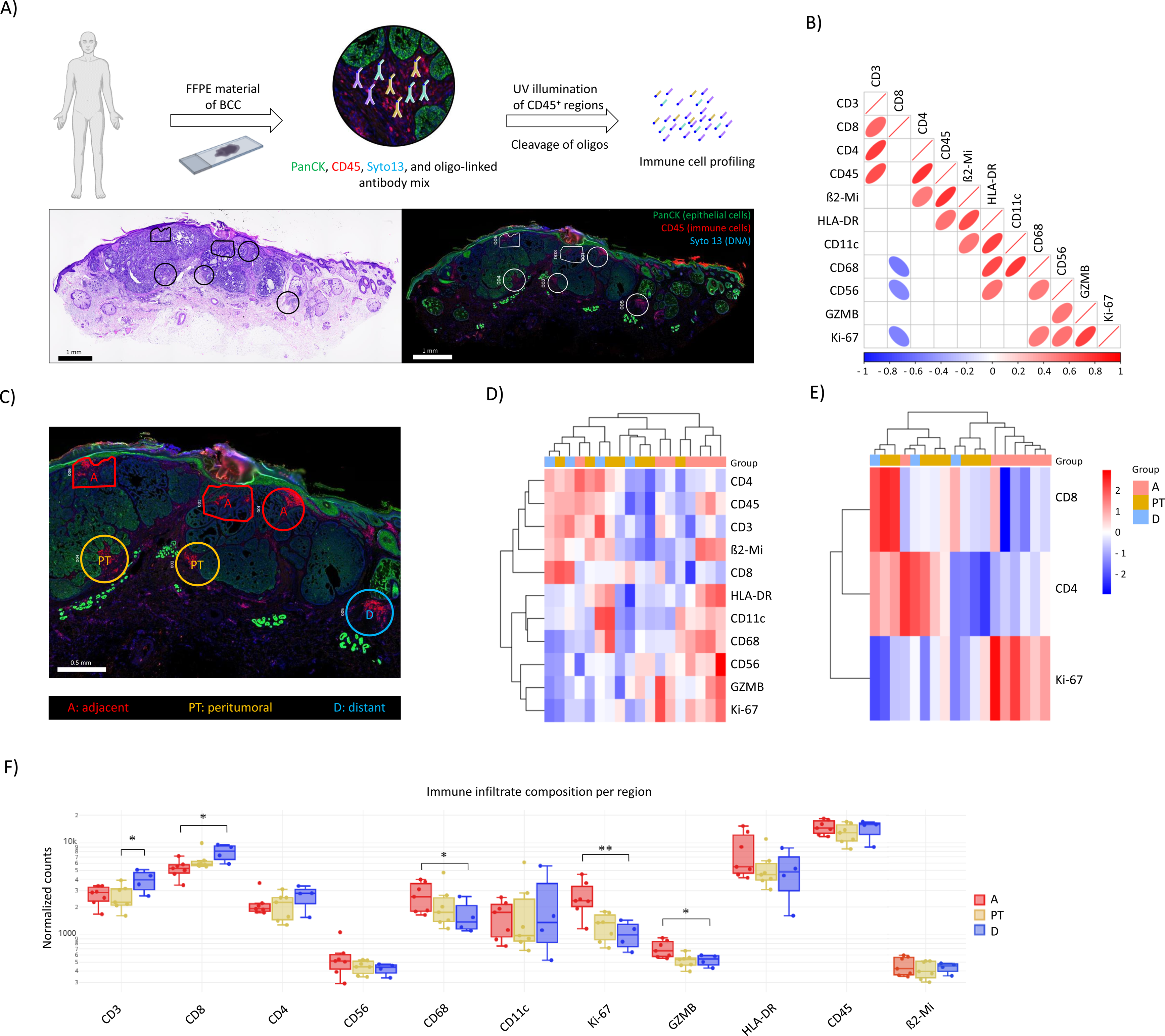
Spatial immune cell profiling of BCC and exclusion of T cells from tumor tissue. **A)** Top: GeoMx DSP workflow: Patient-derived BCC FFPE slides were stained with PanCK (epithelial cells, green), CD45 (immune cells, red), and Syto13 (DNA, blue) for structural guidance and an immune-cell profiling antibody-oligo mix. UV illumination of CD45^+^ regions results in specific cleavage of oligos, which are quantified using the nCounter system. Bottom left: H&E staining of a human BCC biopsy. Scale bar 1 mm. Black regions show different areas within the TME harboring CD45^+^ immune cells selected for harvesting and subsequent analysis. Bottom right: IF staining of PanCK, CD45, and Syto13 of a consecutive section. Scale bar 1 mm. White regions correspond to the selected and harvested CD45^+^ immune cell areas. **B)** Pearson correlation of protein marker expression throughout all analyzed regions. Red: positive correlation, blue: negative correlation. Steepness of elliptical shape corresponds to strength of correlation. Only significant results are shown (p<0.05). **C)** Representative depiction of adjacent (A, red), peritumoral (PT, orange) and distant (D, blue) immune cell infiltrates in the BCC microenvironment. Marked regions correspond to the selected and harvested CD45^+^ immune cell areas. Scale bar 0.5 mm. **D)** Heatmap showing hierarchically clustered expression of marker proteins relative to their own average expression. Colors showing abundance of expression (red higher, blue lower than average) and overlap with A, PT, and D regions. **E)** Hierarchical clustering of T cell marker CD8 and CD4 as well as the activation marker Ki-67 expression (red higher, blue lower than average) and overlap with A, PT, and D regions. **F)** Comparison between marker expression per region category (A, PT, D). N = 7 (A), N = 7 (PT), N = 4 (D). Statistical test: linear mixed model.

Correlation analysis across all 18 regions of interest (ROIs) revealed that areas with high levels of immune activation and proliferation markers GZMB, HLA-DR and Ki-67 predominantly contained CD68^+^ and CD56^+^ cells, most likely representing macrophages and natural killer (NK) cells, respectively. In contrast, CD3^+^ T cell populations, especially the subtype population of CD8^+^ T cells, showed a striking negative correlation with CD68^+^ and CD56^+^ cells, suggesting that T cells mainly localize to different regions within the TME. Notably, regions high in CD8^+^ or CD4^+^ T cells were low in the proliferation marker Ki-67, suggesting that T cells in the BCC tumor niche may be suppressed and are not actively participating in the immune infiltrate (Fig. 3B, suppl. Fig. S3B). Although the panel included the immune checkpoint molecules CTLA-4 and PD-1, neither of them showed a signal above the isotype controls, leaving uncertainty as to whether these T cells may be inactive or exhausted. However, the spatial data suggest that immune cells entering the skin niche segregate and distribute differently in BCC and that the distinct regions differ in terms of immune cell activity.

To determine whether the spatial distribution of immune cell types and activity markers was restricted to specific regions within the tumor or throughout the TME, we selected three distinct groups of ROIs based on the localization of immune cells relative to the primary tumor nodule(s). These groups were termed adjacent (A; red) for immune cells adjacent to the primary tumor site, peritumoral (PT; orange) for immune cells in the tumor vicinity, and distant (D; blue) for immune cells excluded from the tumor tissue (Fig. 3C, suppl. Fig. S3C). Unsupervised hierarchical clustering revealed that immune cells indeed mapped to specific regions within the TME. Specifically, regions with low T cell abundance but high counts of active CD68^+^ and CD56^+^ cells were identified as adjacent regions, suggesting that spatial proximity to the tumor and the depth of penetration predict infiltrate composition (Fig. 3D). To specifically examine the spatial distribution and activity of T cells, we performed a re-clustering of the regions based on CD8 and CD4 T cell markers and Ki-67 as a proxy for activation. We found that both T cell abundance and activity were highly dependent on the proximity to the tumor (Fig. 3E). Comparing distant with adjacent and peritumoral regions, T cells were significantly enriched in distant regions and less likely to be found in closer proximity to the tumor (Fig. 3F). These findings are consistent with our single-cell and 3D OTC analysis and support a model in which HH/GLI and Hippo/YAP crosstalk contributes to a T cell-excluded TME in BCC via disruption of chemotactic signals involved in T cell infiltration and migration.

### Oncogenic GLI and GLI-YAP activation in epidermal cells represses epidermal T cell infiltration

Because human BCC have a complex mutational landscape and multiple driver mutations that potentially affect T cell infiltration patterns independently of GLI-YAP interactions, we used 3D OTCs of G^act^, Y^act^, and GY^act^ to study T cell migration and infiltration behavior specifically in the presence of epidermal HH/GLI, Hippo/YAP, or combined activities. To this end, we introduced human T cells into the dermal layer and observed chemokine secretion of and T cell infiltration into the genetically engineered epidermis. Skin-tropic memory CD4^+^ and CD8^+^ T cells expressing CLA, a P-selectin ligand required for homing to the skin, were isolated from PBMCs of healthy donors (hPBMCs) and mixed in a ratio similar to that of physiological skin (4:1, CD4^+^:CD8^+^)^42^ and seeded together with fibroblasts and fibrinogen to form the dermal layer of a 3D OTC. Control, G^act^, Y^act^, or GY^act^ keratinocytes were then added on top and cultured at the air-liquid interface for 14 days (Fig. 4A).

**Figure 4:**
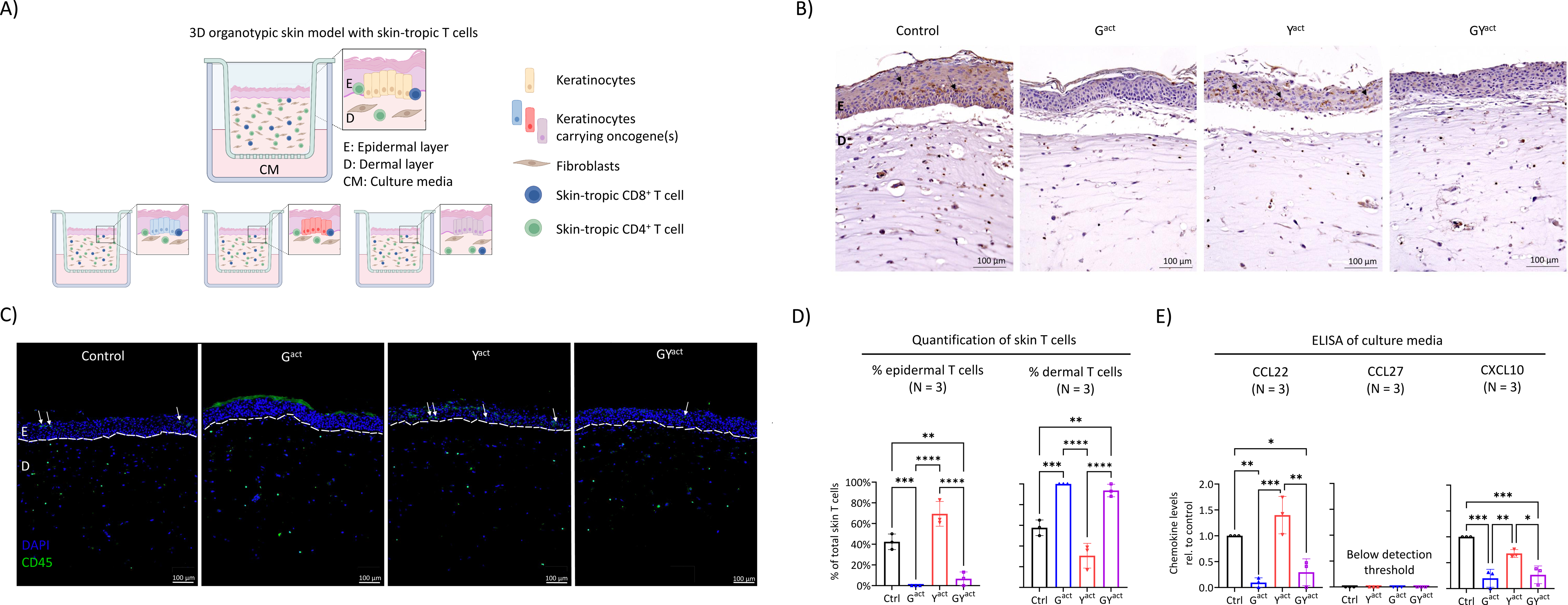
GLI-YAP signal interaction represses epidermal T cell infiltration *in vitro*. **A)** Illustration of the 3D OTC model supplemented with human skin-tropic (CLA^+^) memory (CD45RA^-^) CD4^+^ and CD8^+^ T cells (ratio 4:1) that were isolated by fluorescence-activated cell sorting (FACS) from hPBMCs of healthy donors. The dermal layer consists of fibrinogen, human fibroblasts, and human T cells. Human keratinocytes overexpressing oncogenes were placed on top to grow 3D OTCs under air-liquid interface conditions for 14 days. Data include experiments with hPBMCs of two independent donors. **B)** Representative immunohistochemistry stainings of control, G^act^, Y^act^, GY^act^ 3D OTCs containing human T cells (anti-CD45, brown). Nuclei were counter-stained using hematoxylin (blue). Black arrows indicate epidermal T cells. E: Epidermis, D: Dermis. Scale bars 100 µm. **C)** Representative immunofluorescence stainings of control, G^act^, Y^act^, GY^act^ 3D OTCs containing T cells (anti-CD45, FITC, green). Nuclei were stained using DAPI (blue). White arrows indicate epidermal T cells. The dotted line marks dermal-epidermal junction. E: Epidermis, D: Dermis. Scale bars 100 µm. **D)** Quantification of epidermal (left) and dermal (right) CD45^+^ T cells relative to the total number of T cells per analyzed skin area. N = 3. Statistics: One-way ANOVA. *p<0.05; **p<0.01; ***p<0.001; ****p<0.0001. **E)** ELISA of 3D OTC media for chemokine levels of CCL22, CCL27 and CXCL10. N = 3. Statistics: One-way ANOVA. *p<0.05; **p<0.01; ***p<0.001.

Next, we examined the T cell distribution pattern using human CD45 staining and confirmed the presence of epidermal T cells in both control and Y^act^ 3D OTCs. In stark contrast, G^act^ and GY^act^ suppressed epidermal T cell infiltration (Fig. 4B, suppl. Fig. S4A). We also quantified the relative numbers of epidermal and dermal T cells by immunofluorescence staining for human CD45 coupled with automated cell counting and confirmed that T cells in G^act^ and GY^act^ 3D OTCs remained predominantly in the dermis and barely infiltrated the epidermal layer (Fig. 4C-D). These data are consistent with our analysis of human BCC and suggest that constitutive GLI-YAP activation in the epithelial tumor compartment attenuates T cell infiltration into the tumor tissue. We found that GLI activation alone resulted in efficient exclusion of T cells from the epidermis, suggesting a dominant role of GLI in this process. However, this may also be due to the generally low number of T cells infiltrating the epidermis in 3D OTCs, thereby masking combination effects.

To rule out potential negative effects on T cell viability that might skew their ability to migrate, we also examined the viability of T cells in the entire 3D OTC by flow cytometry. Importantly, T cell viability was comparable in all settings and independent of the epidermal genotype throughout the entire 14-day co-culture period. Furthermore, the ratio of CD4^+^ to CD8^+^ T cells remained constant at the initial seeding ratio of 4:1 and was unaffected by oncogene overexpression in the epidermis (suppl. Fig. 4B). We next examined the surface expression of tissue residency and activation markers (CD69, PD-1) and found that, in agreement with previously published data,^43–45,47^ CD4^+^ and CD8^+^ T cells strongly upregulated CD69 and to a lesser extent PD-1. However, this upregulation was independent of G^act^ or Y^act^ overexpression (suppl. Fig. S4C-D).

To investigate whether the suppression of chemo-attractant factors by GLI and YAP may be responsible for the reduced epidermal infiltration by T cells, we determined the levels of the major T cell chemo-attractants CCL22, CCL27, and CXCL10, which are suppressed in human BCC with high levels of HH signaling (see Fig. 2C), in the media of 3D OTCs by ELISA. As shown in Figure 4E, we detected high levels of CCL22 and CXCL10 in control and Y^act^ 3D OTCs, whereas CCL22 and CXCL10 levels were dramatically reduced in GY^act^ skin. Due to the short-range signaling properties of CCL27, this factor could not be detected in the culture medium, which is located far from the epidermis (Fig. 4E)^33, 46^.

Together with our data on chemokine expression in 3D OTCs and human BCC, these data support a model in which GLI-YAP activity in epidermal cells interferes with epidermal T cell infiltration by downregulating the expression of critical T cell chemo-attractant signals.

### GLI-YAP co-activation attenuates T cell infiltration into the human skin niche and promotes spatial T cell exclusion

To study the actual *de novo* recruitment of T cells into engineered human BCC-like skin over a longer distance and time frame *in vivo*, we used PBMC-humanized immunodeficient NOD- *scid IL2r* ^null^ (NSG) mice bearing dorsal *in situ* generated human skin equivalents consisting of human fibroblasts and keratinocytes^47, 70^. Four weeks after initial skin engraftment and complete wound closure, we transferred hPBMCs into engineered skin-bearing mice, which has been shown to allow for efficient and selective long-term engraftment of human T cells^47^. Four weeks after injection of hPBMCs, we analyzed the spleen and blood for T cell engraftment by flow cytometry. In parallel, we processed FFPE samples of whole human skin equivalents from engrafted mice and examined T cell *de novo* infiltration and their spatial distribution within the human skin equivalent by immunofluorescence (Fig. 5A).

**Figure 5:**
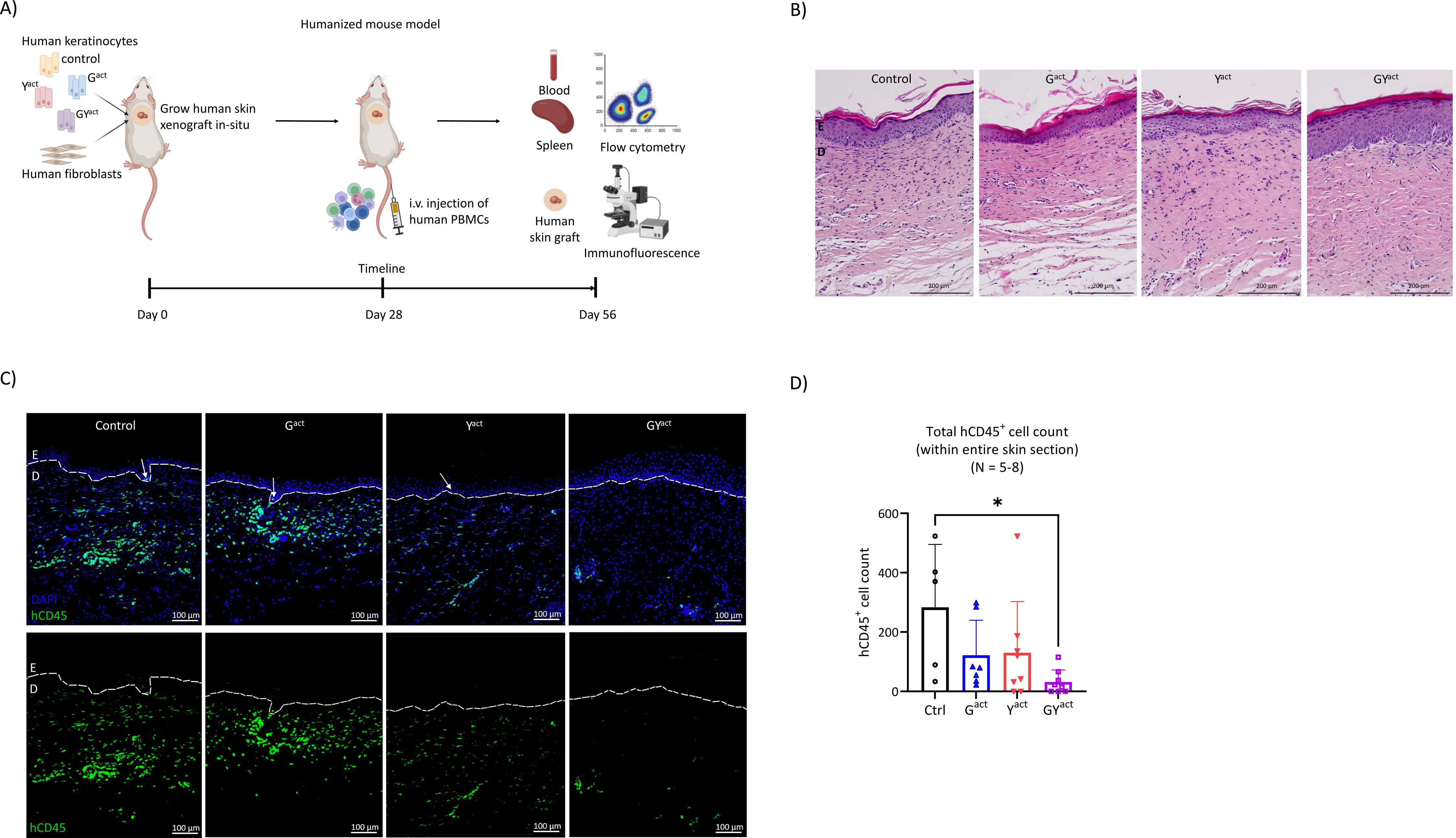
GLI-YAP interaction impairs human T cell infiltration of humanized BCC-like skin *in vivo*. **A)** Illustration of PBMC-humanized immunodeficient NOD-*scid IL2r* ^null^ (NSG) mice. Keratinocytes overexpressing oncogenes together with fibroblasts form a human skin xenograft *in situ* (day 0). After successful engraftment and wound healing, hPBMCs were i.v. injected to complement the hematopoietic system with human immune cells (day 28). Mice were sacrificed 4 weeks later (day 56), spleen and blood were harvested and analyzed via flow cytometry and human skin xenografts were cut out and used for histological analysis. Data include three independent experiments with hPBMCs of three independent donors. **B)** Representative H&E stainings of control, G^act^, Y^act^, GY^act^ 8-week-old human xenograft skins. E: Epidermis, D: Dermis. Scale bars 200 µm. **C)** Representative immunofluorescence images of hCD45^+^ cells (FITC, green) infiltrating control, G^act^, Y^act^, GY^act^ human skin xenografts. Nuclei stained using DAPI (blue). White arrows indicate hCD45^+^ cells infiltrating the epidermal layer. The dotted line represents the dermal-epidermal junction. E: Epidermis, D: Dermis. Scale bars 100 µm. **D)** Absolute number of hCD45^+^ cells per analyzed skin section. Y-axis: hCD45^+^ cell count. N = 5-8. Statistics: Kruskal-Wallis-Test. *p<0.05.

Analyzing the morphology and histology of human skin xenografts, we observed oncogene-dependent alterations of the epidermal layer, especially in GY^act^ skin with hyperproliferation and epithelial downgrowth like 3D OTCs (Fig. 5B). To evaluate and quantify comparable immune cell engraftment efficiencies, we analyzed the spleen and blood of hPBMC-injected mice by flow cytometry. In agreement with previous studies, human immune cells in spleen and blood were composed primarily of CD3^+^ T cells in all mice^44, 47^. As expected, the engraftment of human skin equivalents did not affect the overall engraftment efficiencies and patterns of human immune cells or the ratio of CD4^+^ to CD8^+^ T cells in the spleen or blood (suppl. Fig. S5A-B). Importantly, we found that both T cell subpopulations in spleen and blood expressed surface CLA, a critical marker for skin-homing T cells (suppl. Fig. S5C)^47, 48^. In addition, we observed that T cell numbers in spleen and blood were highly correlated on a per-sample basis (R = 0.66, ***p<0.0095), whereas T cell numbers in human skin xenografts did not correlate with numbers in spleen or blood, indicating specific T cell homing to human skin grafts as previously published for this humanized mouse model (suppl. Fig. S5D)^47^.

To investigate whether T cell infiltration and spatial distribution within human skin grafts was affected by GLI-YAP expression in the human epidermis, we quantified the number of skin-infiltrating hCD45^+^ cells based on immunofluorescence imaging of human skin xenografts (Fig. 5C). For G^act^ and Y^act^ skins, we observed slightly reduced numbers of T cells throughout the human skin graft, whereas human skin grafts with GY^act^ epidermis showed significantly reduced T cell infiltration (Fig. 5D). This suggests that the GLI-YAP interaction inhibits *de novo* T cell recruitment to the skin and thus may spatially exclude T cells from the tumor tissue to facilitate immune evasion.

## Discussion

The study investigates the molecular and cellular effects of combined oncogenic HH/GLI and Hippo/YAP signaling in human epidermis, with a particular focus on chemokine suppression and attenuated T cell infiltration in the TME of BCC. Using 3D human organotypic skin models and humanized mouse models, our research reveals GLI-YAP-mediated perturbations in chemokine networks leading to reduced T cell recruitment and infiltration into tumor tissues.

Examination of bulk and single-cell RNA-seq data from human BCC biopsies confirmed these findings, demonstrating that oncogenic GLI-YAP signaling correlates with low levels of T cell chemo-attractants in human BCC. Notably, spatial profiling of immune cell types in the BCC TME revealed a distinct distribution, with non-proliferative T cells localizing away from primary tumor nodules and proliferative innate immune cells accumulating in adjacent regions.

Furthermore, CD68^+^ innate immune cells (macrophages) mapped primarily to adjacent regions, whereas CD56^+^ innate immune cells (NK cells) were distributed throughout the TME and correlated strongly with activation markers and macrophages. These data suggest that macrophages and NK cells may play a more active role in anti-tumor immunity in BCC, while T cell-mediated responses are impaired due to spatial exclusion. Previous studies have found reduced expression of MHC I molecules in BCC, suggesting enhanced recognition by NK cells, while T cell recognition is further restricted^49, 50^. Alternatively, macrophages could also promote tumorigenesis. In this context, it would be important to characterize the distribution of pro-tumorigenic M2 versus anti-tumorigenic M1 phenotype of macrophages in the TME of BCC in future studies.

Bulk RNA-seq data from primary BCC revealed significantly reduced expression of several T cell chemo-attractants compared to normal skin, including CCL17, CCL22, and CCL27. These chemokines play a critical role in the recruitment of skin-tropic CLA^+^ T cells. NK cells and macrophages respond to a different subset of chemo-attractants, and typical NK cell or macrophage chemo-attractants were not altered in BCC^33, 51–55^. This suggests that the chemokine expression profile of GLI-YAP positive BCC is detrimental for T cell infiltration, while NK cell and macrophage chemotaxis remains largely unaffected, which is consistent with our spatial immune profiling data of BCC.

The study also observed significant patient heterogeneity in chemokine and BCC biomarker expression. Chemokine levels were reduced primarily in patients with high tumor biomarker levels. Single-cell RNA-seq data implicated the BCC driver pathways HH/GLI and Hippo/YAP as regulators of chemokine expression, which was mechanistically confirmed in a 3D OTC model. The interaction of both pathways suppressed chemokine signaling in keratinocytes, including the expression of T cell chemokines CCL22 and CCL27. HH/GLI is essential for BCC growth and is mutated in 90% of patients, with small molecule inhibitors such as vismodegib being highly effective, although resistance is frequently observed^1, 23, 56,57^. Hippo/YAP is mutated in 50% of patients, but activity is reported in the majority of BCC, possibly due to Hippo-independent activation^1, 19^. YAP is mandatory for tumor initiation and blocking of cell differentiation, and YAP target gene expression is stronger in tumors with Hippo pathway mutations, suggesting a conserved immunosuppressive phenotype due to GLI-YAP interaction^1, 58, 59^.

Functional studies in 3D OTCs and human skin xenografts revealed that GLI-YAP-mediated suppression of long-range chemokine signaling is associated with T cell exclusion. By placing skin-tropic T cells near GLI or GLI-YAP oncogene-overexpressing keratinocytes in 3D OTCs, almost complete absence of epidermal T cell infiltration was observed. However, this model does not fully reflect the *in vivo* situation where T cells are recruited *de novo* from secondary lymphoid organs. Using a humanized NSG mouse model, the study found that the combination of GLI-YAP, but not GLI alone, attenuated *de novo* T cell recruitment and promoted T cell exclusion from the skin niche. This suggests that T cell exclusion in BCC is established at the level of extravasation, involving CCL22 and CCL27 via CCR4 and CCR10 interaction.

Spatial data of human BCC show that T cells are present within the tumor skin niche but localize to tumor-distant areas near healthy tissue. BCC is surrounded by healthy skin, including keratinocytes that do not carry HH/GLI and Hippo/YAP mutations and can respond to inflammatory cues by producing chemokines such as CCL22 and CCL27. This may initially recruit T cells to the skin niche, but these T cells are unable to enter the TME due to disrupted chemokine networks caused by GLI-YAP. The study demonstrates that GLI-YAP inhibits long-range chemokine signaling, leading to imbalances in chemokine networks in human BCC that prevent T cells from entering tumor areas.

The results suggest that *de novo* infiltration of T cells and their ability to reach the primary tumor in BCC is regulated by oncogenic HH/GLI and Hippo/YAP signaling. Information about the tumor oncogenome, particularly mutational activation of HH/GLI and Hippo/YAP, has potential value in predicting response to immunotherapy in BCC patients. In summary, these new insights into the molecular control of the tumor immune microenvironment of BCC may provide the basis for improved combination therapies involving treatments that promote T cell recruitment, including inhibition of HH/GLI and Hippo/YAP signaling, together with immune checkpoint blockers to reinitiate efficient anti-tumor immune responses.

## Materials and Methods

### Ethics

This study was conducted according to the approved guidelines of the World Medical Association’s Declaration of Helsinki and the guidelines of the Ethics Committee of the Province of Salzburg and the Austrian national regulations.

### GeoMx digital spatial profiling

Human BCC samples were obtained from the Department of Dermatology, Paracelsus Medical University/Salzburger Landeskliniken (SALK) from patients undergoing surgical excision and prepared as 5 µm thick FFPE sections. Section preparations included manual deparaffinization and rehydration (Xylol, 100% EtOH, 95% EtOH, Water), antigen retrieval (Citrate pH6, pressure cooker), blocking with Nanostring Buffer W, and staining with immunofluorescence biomarkers (PanCK-AF594, CD45-AF532, DNA Syto13 (Nanostring)) and antibodies linked to photo-cleavable DNA tags (Human protein core immune cell profiling panel (Nanostring): Beta-2-microglobulin, CD11c, CD3, CD4, CD8, CD20, CD45, CD56, CD68, CTLA4, Fibronectin, HLA-DR, GZMB, Ki-67, PanCK, PD-1, PD-L1, SMA, and isotype/housekeeping protein controls). Slides were scanned with the GeoMx DSP system to create digital images highlighting tissue features. Regions of interest (ROIs) were selected based on CD45 expression and tissue characteristics. CD45-positive areas were exposed to UV light to cleave DNA tags and the oligos were quantified using the nCounter system. Background normalization was performed with IgG negative controls and used to identify and exclude non-expressed targets. Normalized counts were mapped, analyzed, and plotted using GeoMx DSP analysis suite (version 2.4.0.421), SRplot^60^ or GraphPad Prism (version 9.0).

### In silico analysis of bulk & single cell RNA-seq data

Bulk RNA-seq data of 24 human normal skin and 59 BCC biopsies were provided by Bonilla et al. (2016)^1^ and re-analyzed for DEGs based on initial classification (“Tumor” vs “Normal”) via limma-voom in R. Volcano plot was generated using VolcaNoseR^61^ displaying genes with a Log_2_ fold-change of > 1 or < −1 and q_val_ < 0.05. Hierarchical clustering of the selected list of DEGs was done using R. Single cell RNA-seq data was obtained via Gene Expression Omnibus (accession number: GSE123814) published by Yost et al. (2019)^41^ and patients harboring nodular or infiltrative BCC (N = 7) pre-anti-PD-1 treatment were selected for further analyses. Original clustering of cell types was used and further aggregated to 10 cell types for analysis. Tumor populations 1 and 2 were combined to derive oncogenic signatures. Pseudo-bulk expression values were obtained by taking mean across all cells per cell type and sample. Oncogenic signatures for HH/GLI (PTCH1, PTCH2, GLI1, GLI2) and Hippo/YAP (CYR61, CTGF, CRIM1, TGFB2, AXL) activity were derived from the Molecular Signatures Database and manual curation according to tissue-specific expression of target genes in skin^18, 41, 62–65^. Pearson correlation was calculated using SRplot^60^. T cell abundance was calculated relative to the total number of immune cells per biopsy and correlated with the oncogenic signature strength of HH/GLI or Hippo/YAP in corresponding tumor cells. Benjamini-Hochberg method of multiple hypothesis testing was used to calculate p_adj_ (q_val_) from p_val_.

### Cell Culture and retroviral transduction

N/TERT-1 human keratinocyte cells (kindly provided by James G. Rheinwald^31^) were certified by DSMZ via fingerprinting and cultured under sub-confluent conditions at 37 °C and 5% CO_2_ in Epilife media (ThermoFisher) supplemented with 5 mL Human Keratinocyte Growth Supplement, 100 U/mL penicillin and 100 mg/mL streptomycin. Fibroblasts were isolated from normal human skin and immortalized using human papilloma virus type oncogenes E6/E7 as previously described^66^ and cultured at 37 °C and 5% CO_2_ in DMEM (Sigma) supplemented with 10% fetal bovine serum (FBS) (Sigma), 100 U/mL penicillin and 100 mg/mL streptomycin (Sigma). Phoenix-Ampho cells (ATCC) were cultured in DMEM supplemented with 10% FBS, 2 mM L-GLUT (Sigma), 0.1 mM NEAA (Sigma), 100 U/mL penicillin and 100 mg/mL streptomycin.

Both the retroviral overexpression construct pBABE puromycin FLAG-YAPS127A (Biocat) and the in-house established pMSCV hygromycin HA-ΔNGLI2 were sequence-verified^65^. Retroviral particles were produced by introducing transfer-plasmids into Phoenix-Ampho cells. Supernatant enriched for virions was harvested, screened for particle density via q- PCR (Retrovirus Titration Kit, Abcam), and used to transduce N/TERT-1 keratinocytes with a MOI of 100 via spinfection using Polybrene (Sigma) as transfection reagent. Successfully transduced keratinocytes were selected with 0.5 µg/mL Puromycin (Sigma) for 5 days and 20 µg/mL Hygromycin (Sigma) for 7 days.

### Western blot analysis

One well of a 6-well plate containing a confluent layer of N/TERT-1 cells overexpressing oncogenes or vector controls was harvested in 1x Laemmli buffer supplemented with phosphatase and protease inhibitors (cOmplete Mini, Sigma)^67^. Proteins were separated by SDS-PAGE and blotted onto nitrocellulose membranes (Life Technologies). Antibodies used to verify oncoprotein expression included primary antibodies anti-HA-tag (CST, rabbit, 1:1000), anti-FLAG-tag (CST, rabbit, 1:1000), anti-total-ERK1/2 (CST, rabbit, 1:1000) diluted in PBS containing 1% BSA (Sigma); and secondary antibody anti-rabbit (CST, 1:3000) diluted in PBS containing 4% skim-milk powder (Merck) and 0.2% Tween 20 (Sigma).

### 3D organotypic skin cultures (OTCs)

The dermal matrix of 3D OTCs was prepared in 12-well transwell inserts with 0.4 µm pore size (GBO) and consisted of 2.7*10^5^ fibroblasts in DMEM supplemented with 10% FBS, 100 U/mL penicillin and 100 mg/mL streptomycin, fibrinogen (final conc. of 7.35 mg/mL, from human plasma, Sigma), thrombin (from human plasma, Sigma) and aprotinin (from bovine lung, Sigma). For the establishment of 3D OTCs with T cells, human skin-tropic (CLA^+^) memory (CD45RA^-^) CD4^+^ and CD8^+^ T cells were isolated from hPBMCs, mixed in a 4:1 ratio (2.4*10^5^ CD4^+^ and 0.6*10^5^ CD8^+^), added to the dermal matrix together with 2.7*10^5^ fibroblasts and incubated at 37 °C and 5% CO_2_ for 1 hour to complete matrix stiffening. Subsequently, 2.7*10^5^ N/TERT-1 keratinocytes overexpressing oncogenes or vector controls were seeded as monolayer directly on top of the matrix in Greens keratinocyte seeding media consisting of 260 mL DMEM, 120 mL F12 nutrient mix (ThermoFisher) supplemented with 10% heat-inactivated FBS, 8 mL 200 mM L-GLUT, 4 mL 100 mM sodium-pyruvate, 0.14 mM adenine (Sigma), 0.5 µg/mL hydrocortisone (Sigma), 5 µg/mL insulin (Sigma), 4.23 ng/mL cholera toxin (Sigma), 1.37 ng/mL triiodothyronine (Sigma), 100 U/mL penicillin and 100 mg/mL streptomycin. After 24 hours of culture, the media was removed and skins were cultured in 12-well deep-well plates (GBO) under air-liquid interface conditions in Greens^+^ keratinocyte proliferation media (Greens media supplemented with 10 ng/mL human EGF (Sigma)). Media was exchanged every other day and from day 7 onwards Vitamin C (Sigma) was added daily (final conc. 50 µg/mL, Sigma) until harvest of skins on day 14.

### RNA isolation, real-time quantitative polymerase chain reaction (q-PCR), RNA-sequencing and bioinformatic analysis

The epidermis was separated from the dermal layer via digestion in Dispase II (final conc. 2.7 U/mL in PBS, Roche) for 30 min on ice. The epidermis was subsequently stripped off the dermal layer, washed in PBS and immediately transferred into TRI reagent (Sigma) and minced via Ultra Turrax T8 until completely dissolved. Total RNA was isolated using LiCl precipitation. Quality of RNA was assessed using an Agilent 2100 Bioanalyzer (RIN > 9, 28S/18S ratio > 1 for all samples). For q-PCR, 500 ng of RNA were reverse transcribed into cDNA using M-MLV reverse transcriptase (Promega). q-PCR was performed using GoTaq q- PCR Mastermix (Promega) according to manufacturer’s instructions on a Rotor-Gene Q instrument (Qiagen). RNA-seq, including library preparation, was performed by BGI Tech Solutions on a DNBSEQ platform with a minimum of 20 million read pairs per sample (N = 3 per condition). Raw data processing, including quality control, human genome alignment and gene expression quantification was done by BGI. DEG analysis for all comparisons was done via limma-voom in R. DEGs for all analyses represent genes with a Log_2_ fold-change > 1 or < −1 and q_val_ < 0.05. GO BP and TRRUST enrichment analyses were done using the Enrichr tool^68, 69^. Benjamini-Hochberg method of multiple hypothesis testing was used to calculate p_adj_ (q_val_) from p_val_.

### Human PBMC isolation from whole blood

PBMCs were isolated from fresh buffy coats from healthy, anonymous donors provided by the blood bank Salzburg, Austria. Briefly, PBMCs were isolated by density gradient centrifugation using Histopaque-1077 (Sigma). After erythrocyte lysis using ACK buffer (150 mM ammonium chloride, 10 mM potassium bicarbonate, 0.1 mM EDTA, pH7.4) cells were washed twice with PBS and frozen in RPMI-1640 (Sigma), supplemented with 40% heat-inactivated fetal calf serum (Biowest), 10% dimethyl sulfoxide (Carl Roth GmbH) and 2 mM L-GLUT.

### Cell sorting and flow cytometry

Previously cryopreserved hPBMCs were cultured overnight at 37 °C and 5% CO_2_ in T cell media (RPMI-1640 supplemented with 5% heat-inactivated FBS (Sigma), 100 U/mL penicillin, 100 mg/mL streptomycin, 2 mM L-GLUT, 0.1 mM NEAA, 1 mM Sodium-Pyruvate (Sigma), 10 mM HEPES (Sigma), 50 µM ß-Mercaptoethanol (Lactan)), washed in cold PBS, incubated in FC-blocking mix (PBS containing human anti-CD16/CD32/CD64, 1:100 dilution, BioLegend) for 10 min at 4 °C and subsequently stained for surface markers in antibody mix in PBS for 30 min at 4 °C in the dark. Surface staining antibodies were obtained from BioLegend and diluted in PBS for staining (CLA-FITC clone HECA-452 1:20 dilution; CD45RA-AF700 clone HI100, 1:50 dilution; CD4-PE-Dzz/594 clone RPA-T4, 1:100 dilution; CD8-BV510 clone RPA-T8, 1:100 dilution). Viability dye eFluor780 (1:1000 dilution) was obtained from ThermoFisher. Using a FACS Aria III (BD Biosciences), CLA^+^ CD45RA^-^ CD4^+^ and CD8^+^ T cells were sorted into individual 15 mL tubes containing DMEM supplemented with 10% heat-inactivated FBS, 100 U/mL penicillin and 100 mg/mL streptomycin, washed with PBS and mixed in a 4:1 ratio (CD4^+^ to CD8^+^) in DMEM for subsequent seeding into 3D OTCs.

Heart-drawn blood was collected and immediately mixed with 20 µL 0.1 M EDTA to avoid clotting. Spleens were meshed in PBS. Skins were cut into small pieces and digested in 0.8 mg/mL collagenase IV solution (Worthington) in T cell media for 3 hours at 37 °C and 5% CO_2_. For the last 15 min of incubation, 11.77 U/mL DNAse I solution (Sigma) was added. Cell suspensions from 3D OTCs and spleen were strained through 100 µm cell strainer. Samples were cooled on ice and incubated in surface antibody staining solution in PBS for 30 min at 4 °C in the dark. Blood samples were incubated and washed with RBC Lysis/Fixation Solution (BioLegend) for 4 min at RT. Surface staining antibodies were obtained from BioLegend and diluted in PBS for staining (CLA-FITC clone HECA-452, 1:20 dilution; CD69-PerCP Cy5.5 clone FN50, 1:20 dilution; CD4-PE-Dzz/594 clone RPA-T4, 1:100 dilution; PD-1-PE-Cy7 clone eBioJ105, 1:20 dilution; CD3-BV421 clone UCHT1, 1:100 dilution; CD8-BV786 clone RPA-T8, 1:200 dilution; CD45-Pe594 clone HI30, 1:200 dilution; CD4-PE-Cy5 clone RPA-T4, 1:100 dilution; CD45RA-BV510 clone HI100, 1:100 dilution; CLA-BV605 clone HECA-452, 1:80 dilution; CD8-BV785 clone RPA-T8, 1:100 dilution). Viability dye eFluor780 (1:1000) dilution was obtained from ThermoFisher. All samples were finally washed and solved in PBS for flow cytometric analysis performed on a CytoFLEX S (Beckman Coulter).

### Enzyme-linked Immunosorbent Assay (ELISA)

The entire culture media from day 13-14 of the 3D OTC was transferred into a 15 mL tube, immediately shock-frozen in liquid nitrogen and stored at −80 °C for later analysis. The human CCL22/MDC ELISA kit was purchased from R&D Systems, Biotechne. The human CCL27/CTACK and CXCL10/IP-10 were purchased from Peprotech. The assays were performed on supernatants according to the manufacturer’s instructions and the readout was done on a Tecan microplate reader (Infinite 200 pro).

### Immunofluorescence and immunohistochemistry

Fresh tissue samples were fixed in 4% PFA overnight, embedded in paraffin and sectioned as 5 µm thick slides using a Leica microtome (Leica Biosystems) followed by deparaffinization and rehydration (Xylol, 100% EtOH, 50% EtOH, PBS) and either stained in aqueous solution of H&E (Hematoxylin Gill II, Roth; Eosin G-Solution 0.5%, aqueous, Lactan) for 12 min and 5 min respectively or used for immunofluorescence or immuno-histochemistry. Heat-induced epitope retrieval was done in citrate buffer with pH6 for all stainings. Retrieval was achieved by shortly boiling of the citrate solution in a microwave for three consecutive times, with 10 min of cooling in-between heating cycles. Afterwards, samples were thoroughly washed in PBS and blocked in blocking buffer (1% BSA in PBS) for 1 hour at RT to saturate unspecific binding sites. Immunofluorescence samples were incubated in primary antibody solution (anti-hCD45 rabbit, CST, clone D9M8I, 1:200 in blocking buffer) overnight at 4 °C and subsequently in secondary antibody solution (anti-rabbit IgG, CST, Alexa Fluor 488 conjugate, 1:1000 in blocking buffer) for 1 hour at RT. Stained slides were covered in DAPI mounting media (Abcam). Immunohistochemistry staining procedure included incubation of samples in primary antibody solution (anti-hCD45 rabbit, CST, clone D9M8I, 1:200 in blocking buffer) overnight at 4 °C, blocking of endogenous peroxidase activity in 3% H_2_O_2_ for 10 min at RT, use of an HRP-linked secondary antibody (HRP-linked anti-rabbit, CST, not diluted) for 30 min at RT, incubation of slides with a 3% DAB solution (CST) for 5 min, washing in PBS, hematoxylin counterstaining, dehydration, and coverage of slides in synthetic resin (RotiHistoKitt II, Roche). Imaging was done using a Nikon Ti-2 microscope and the NIS-Elements software (version 5.40.00). For T cell quantification in 3D OTCs and human xenografts, whole skin sections were imaged, and the central human skin graft region was analyzed using the watershed algorithm in the NIS-Elements software followed by automated cell counting of hCD45^+^ cells (FITC signal).

### Human skin xenografts in humanized NSG mice

Immunodeficient NOD-scid IL2rγ^null^ (NOD.Cg-Prkdcscid Il2rgtm1Wjl/SzJ) mice purchased from the Jackson Laboratory were maintained in a specific pathogen-free facility at the Central Animal Facility of the University of Salzburg. The animal studies were approved by the Austrian Federal Ministry of Science, Research and Economy (BMWFW, application no. GZ 2020-0.437.219).

Human engineered skins were generated in NSG mice as described previously^47^ using immortalized human keratinocytes and fibroblasts. Human skin xenograft tissue was generated *in -vivo* in mice by mixing 10^6^ keratinocytes (either control, Y^act^, G^act^, or GY^act^) with 10^6^ fibroblasts in 400 µL MEM (Gibco) containing 1% FBS, 2 mM L-GLUT and 0.1 mM NEAA and placing it in silicone grafting chambers surgically mounted on the back of the mice as previously described^47^. The grafting chambers were removed one week after transplantation and skin wounds were allowed to heal completely within 4 weeks, thereby forming human skin xenografts. Previously cryopreserved hPBMCs were cultured overnight at 37 °C and 5% CO_2_ in T cell media before engraftment. Mice received an i.p. injection of 100 µg anti-Gr1-antibody (clone RB6-8C5, Biozol) in 200 µL PBS the day prior to hPBMC injection to deplete murine granulocytes. Engraftment of 2.5*10^6^ hPBMCs in 200 µL PBS was done via i.v. injection. Mice were sacrificed 4 weeks after hPBMC injection and harvested for blood, spleen, and human xenograft skin analysis.

### Statistical analysis

Statistical analyses were carried out using either the GeoMx DSP analysis suite (version 2.4.0.421, Nanostring) for GeoMx data or the GraphPad Prism (version 9.0). Unless otherwise indicated, data were tested for normality (Shapiro-Wilk test, Kolmogorov-Smirnov test) and either one-way ANOVA or Kruskal-Wallis was used to compare data sets. Graphs depict mean values ± standard deviation (indicated as error bars), and sample numbers as well as p-values are mentioned in the figures. Levels of significance were subdivided into the following categories: *p<0.05; **p<0.01; ***p<0.001; ****p<0.0001. Correlation analyses were calculated by Pearson correlation test using SRplot^60^ or GraphPad Prism. Volcano plots were done using VolcanoseR^61^. Pathway enrichment analyses were done using Enrichr^68, 69^. In cases of multiple hypothesis testing, Benjamini-Hochberg method was used to calculate p_adj_ (q_val_) from p_val_.

## Acknowledgements

The authors are thankful for the help of the animal caretakers and the head of the animal facility, Dr. Angelika Sales for input regarding mouse experiments. We acknowledge the technical support by Sophie Aichberger, Sabine Siller, Anna Eglseer, and Mag. Christian Behensky and the scientific input by Florian Rathje, Daniel Lankes, Lisa Trattner and all other members of the Aberger group for continuous scientific and technical support. We would also like to express special thanks to Dr. Sergey Nikolaev for providing access to raw count data from Bonilla et al. (2016). The following illustrations were created with BioRender.com: Fig. 1A; Fig. 2A, D; Fig. 3A; Fig. 4A; Fig. 5A).

## Author contributions

GS, SRV, PWK, DHH, AS performed all key experiments. GS, SRV, PWK, DPE, DN, JHH, NF, IKG, FA designed experiments, generated and evaluated data and/or provided biological samples and material. MS, NZ, DN, and RG provided technical and intellectual support and/or performed experiments. GS, SRV, PWK, DPE, JHH, NF, IKG, FA analyzed, discussed, and interpreted data. GS, PWK and FA wrote the manuscript. All authors read and approved the final manuscript.

