## supplemental information for "Hedgehog-Hippo pathway interactions promote T cell exclusion from the tumor microenvironment in basal cell carcinoma"

Supplementary figure legends:

**Suppl. Figure S1: A)** Top recurrent pathway mutations in human BCC based on exon and cancer panel sequencing by Bonilla et al. (2016)[1]. **B)** Left: Representative Western Blot of N/TERT-1 overexpressing oncoproteins (top: HA-tagged ΔNGLI2; middle: FLAG-tagged YAP; bottom: total-ERK1/2) after retroviral transduction in 2D culture. Right: quantification of FLAG-tagged $YAP^{S127A}$ and HA-tagged ΔNGLI2 oncoprotein levels in N/TERT-1. N = 3. **C)** Transcript levels of total YAP and total GLI2 in 3D OTC epidermis relative to the housekeeping gene RPLP0 measured via q-PCR. N = 5. Statistics: one-way ANOVA for YAP, Kruskal-Wallis for GLI2. *p<0.05; **p<0.01; ****p<0.0001. **D)** Transcript levels of CCL22 and CCL27 in 3D OTC epidermis relative to the housekeeping gene RPLP0 measured via q-PCR. N = 5. Statistics: one-way ANOVA for CCL22, Kruskal-Wallis for CCL27. *p<0.05; **p<0.01; ***p<0.001; ****p<0.0001.

**Suppl. Figure S2: A)** HH/GLI (PTCH1, PTCH2, GLI1, GLI2) and Hippo/YAP (CYR61, CTGF, CRIM1, TGFB2, AXL) gene signature composition deployed to derive pathway activity in tumor cells. **B)** Relative, mean HH/GLI (top row) and Hippo/YAP (bottom row) gene signature strength in all tumor cells per patient (Pat1-7). **C)** Pearson correlation between HH/GLI and Hippo/YAP gene signature strength in tumor cells of all patients (Pat1-7).

**Suppl. Figure S3: A)** H&E and PanCK, CD45, Syto13 stainings of all human BCC samples (Pat1-3) used for GeoMx DSP. Green: epithelial cells (PanCK); red: immune cells (CD45); blue: DNA (Syto13). Circled regions in white correspond to the selected and harvested $CD45^+$ immune cell areas. Scale bars 1 mm. **B)** Pearson correlation of normalized expression of Ki-67 vs CD8, Ki-67 vs CD4, CD8 vs CD68, CD8 vs CD56, CD4 vs CD68, CD4 vs CD56, CD56 vs CD68, CD56 vs GZMB amongst all 18 ROIs. p-values depicted in graphs. **C)** Regions harboring adjacent (A, red), peritumoral (PT, orange), and distant (D, blue) immune cell infiltrates (Pat1-3). Scale bars as indicated (0.5 mm or 1 mm).

**Suppl. Figure S4: A)** Representative immunohistochemistry stainings of control, $G^{act}$, $Y^{act}$, $GY^{act}$ 3D OTCs containing T cells (upper row) or without T cells (lower row) stained for anti-CD45 (brown). Nuclei were counter-stained using hematoxylin (blue). Black arrows indicate epidermal T cells. E: Epidermis, D: Dermis. Scale bars 100 µm. **B)** T cell viability (left) and the $CD4^+$:$CD8^+$ T cell ratio (right) in 3D OTCs as determined by flow cytometry. N = 2. Statistics: comparisons by Kruskal-Wallis were not significant. **C)** Fluorescence intensity distribution of $CD69^+$ or PD-$1^+$ $CD3^+$ T cells from ingoing hPBMCs (dark grey), control (grey), $Y^{act}$ (red), $G^{act}$ (blue) or $GY^{act}$ (turquoise) 3D OTCs. N = 2. **D)** Percentage of $CD8^+$ (upper row) and $CD4^+$ T cells (lower row) in 3D OTCs that stained positive for surface CD69 or PD-1 expression. N = 2. Statistics: comparisons by Kruskal-Wallis were not significant (not shown).

**Suppl. Figure S5: A-B)** Flow cytometry data of spleen (A) and blood (B) showing percentage of $hCD45^+$ cells relative to all viable cells, percentage of $CD3^+$ cells amongst $hCD45^+$ cells, percentage of $CD4^+$ and $CD8^+$ cells of $CD3^+$ T cells, and the ratio between $CD4^+$:$CD8^+$ T cells. N = 5-8 for spleen, N = 2-4 for blood. Statistics: comparisons by one-way ANOVA were not significant (not shown). **C)** Percentage of skin-tropic memory T cells ($CLA^+$ $CD45RA^-$) of $CD4^+$ and $CD8^+$ T cell subpopulations in spleen (left) and blood (right). N = 5-8 for spleen, N = 2-4 for blood. Statistics: comparisons by one-way ANOVA were not significant (not shown). **D)** Pearson correlation between the percentage of $hCD45^+$ cells in spleen, blood (measured via flow cytometry) and skin (percentage of $hCD45^+$ cells of all $DAPI^+$ cells per analyzed skin section, measured by immunofluorescence imaging) for pooled samples. Red: significantly positively correlated (***p<0.001); grey: not significantly correlated.
