## supplemental figures S1-S5 for "Hedgehog-Hippo pathway interactions promote T cell exclusion from the tumor microenvironment in basal cell carcinoma"

Stockmaier et al., suppl. Figure S1

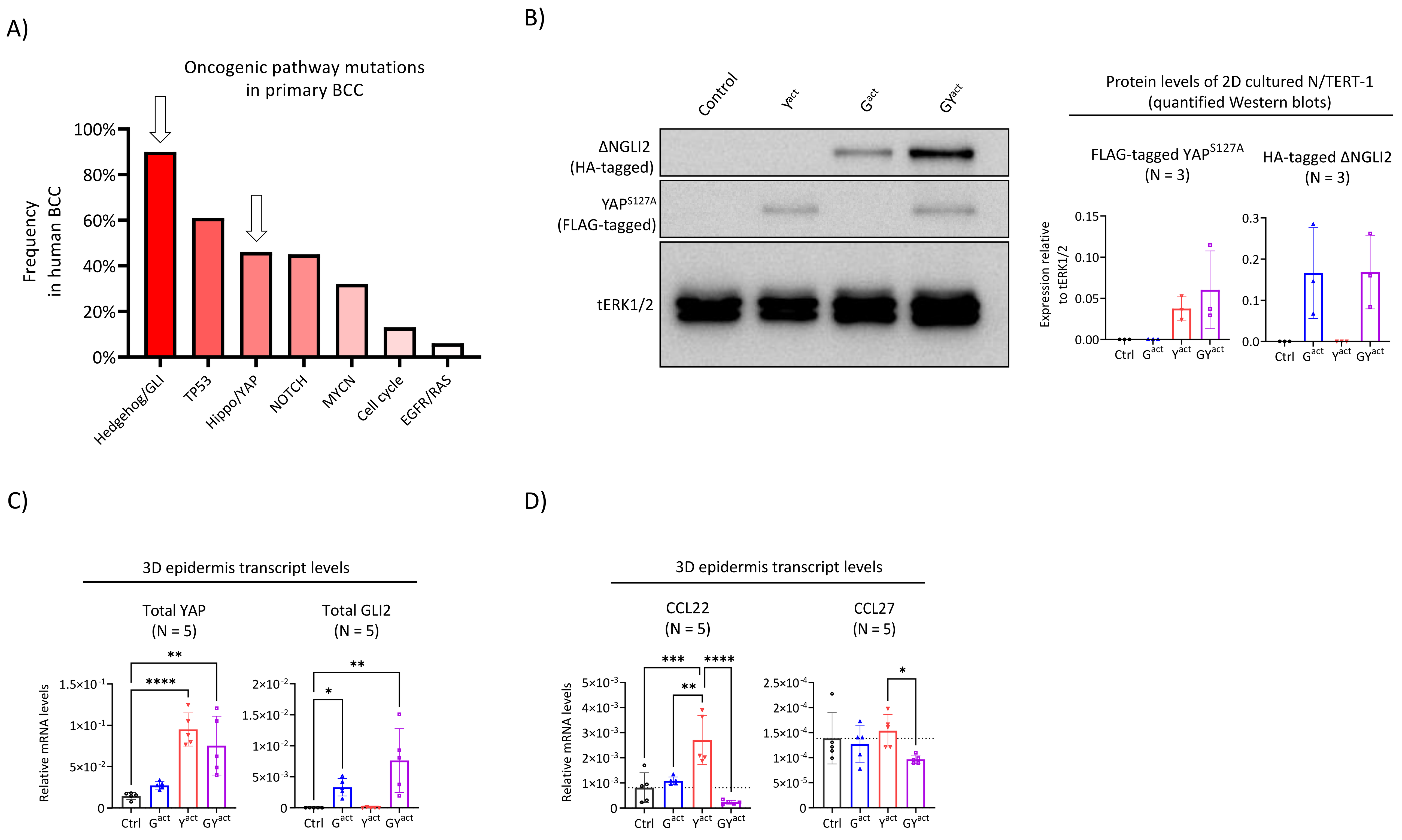

Stockmaier et al., suppl. Figure S2

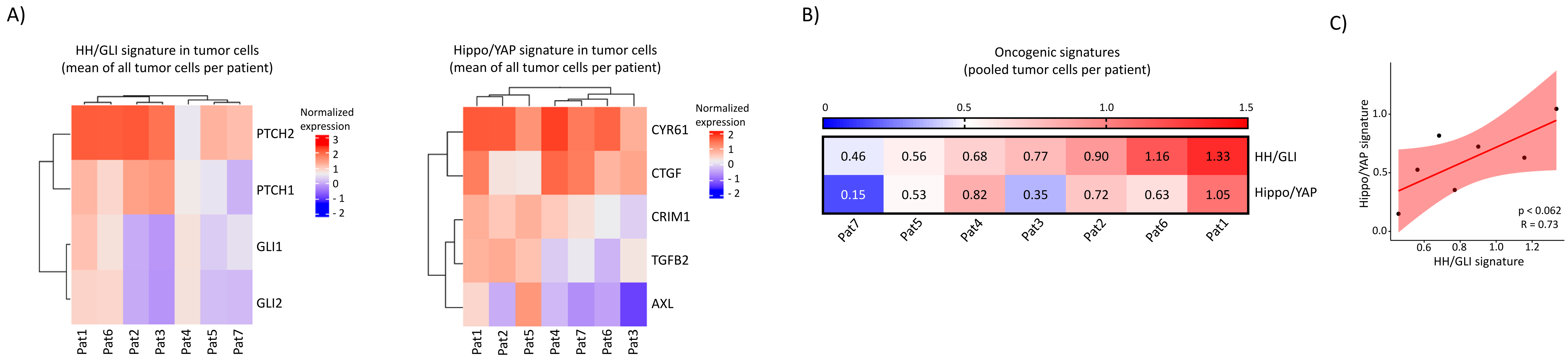

### Stockmaier et al., suppl. Figure S3

A)

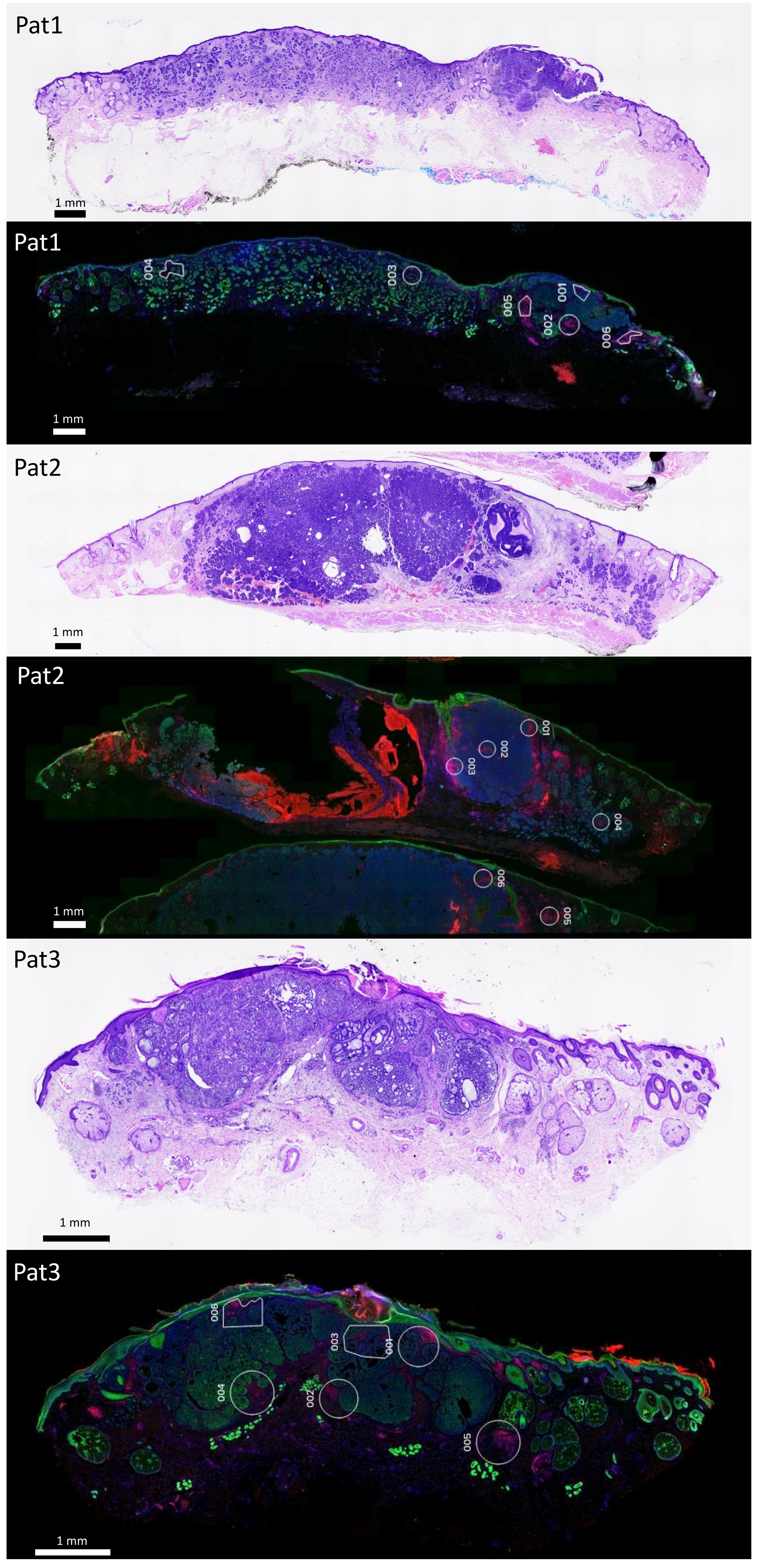


B)

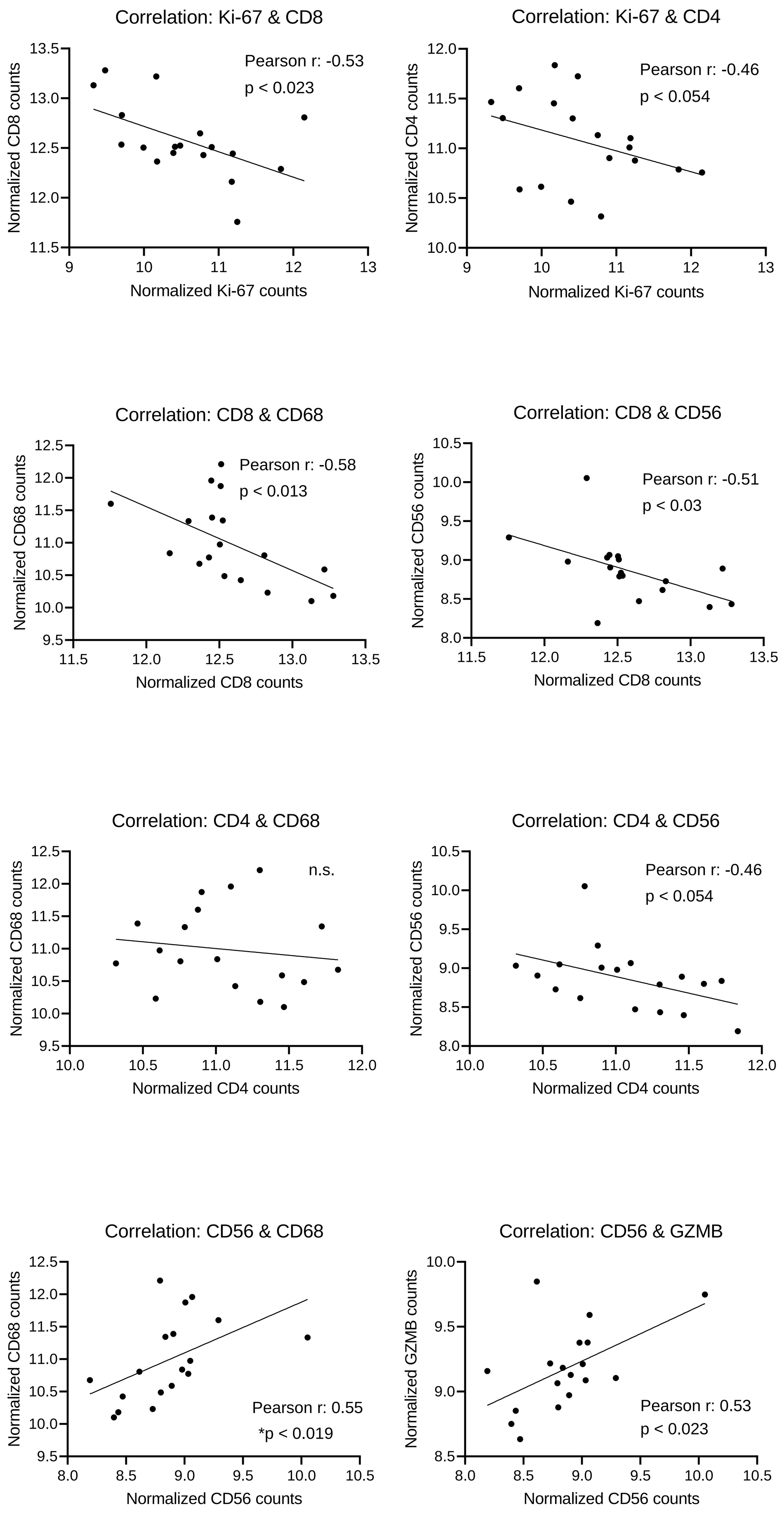


C)

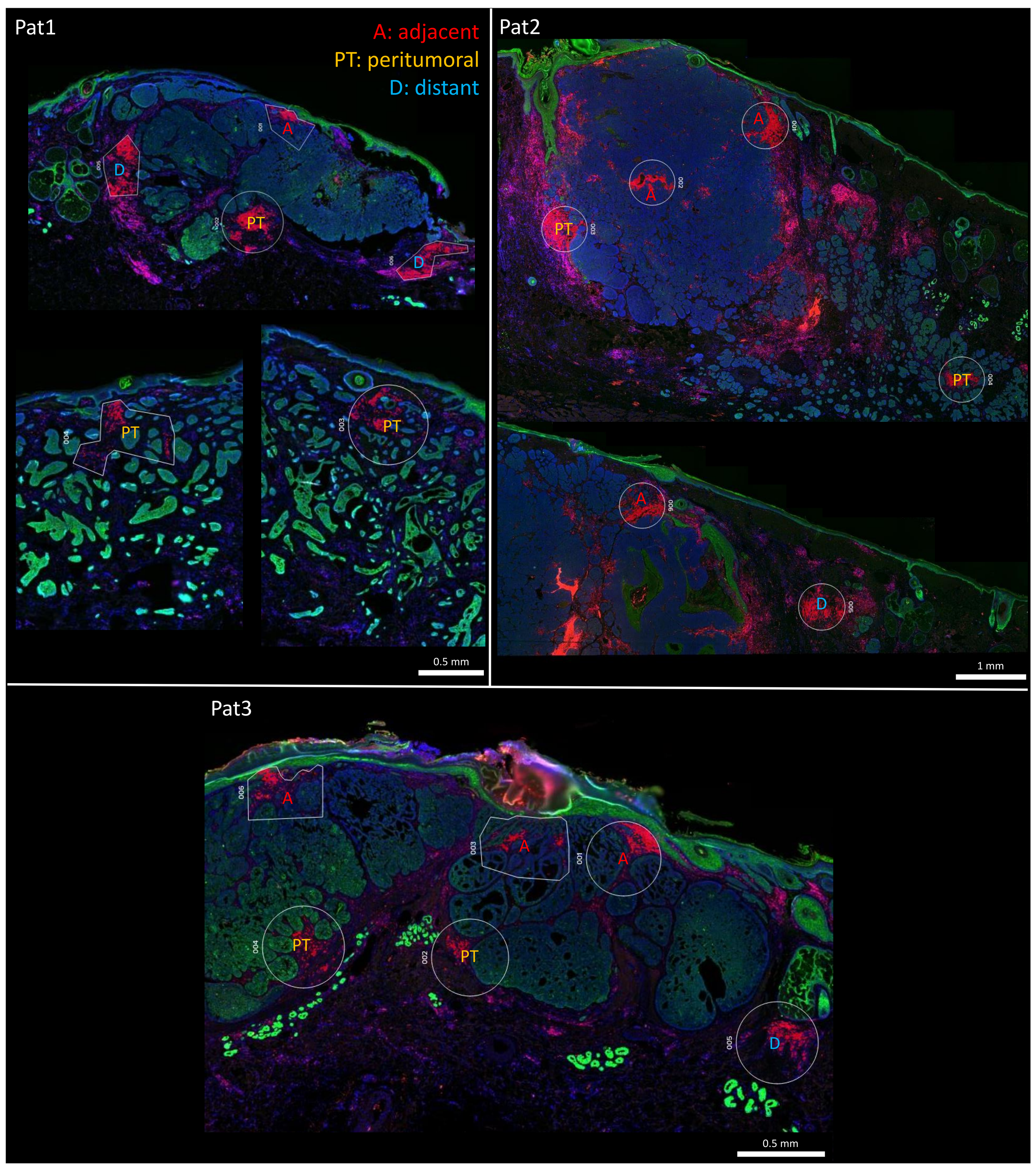

### Stockmaier et al., suppl. Figure S4

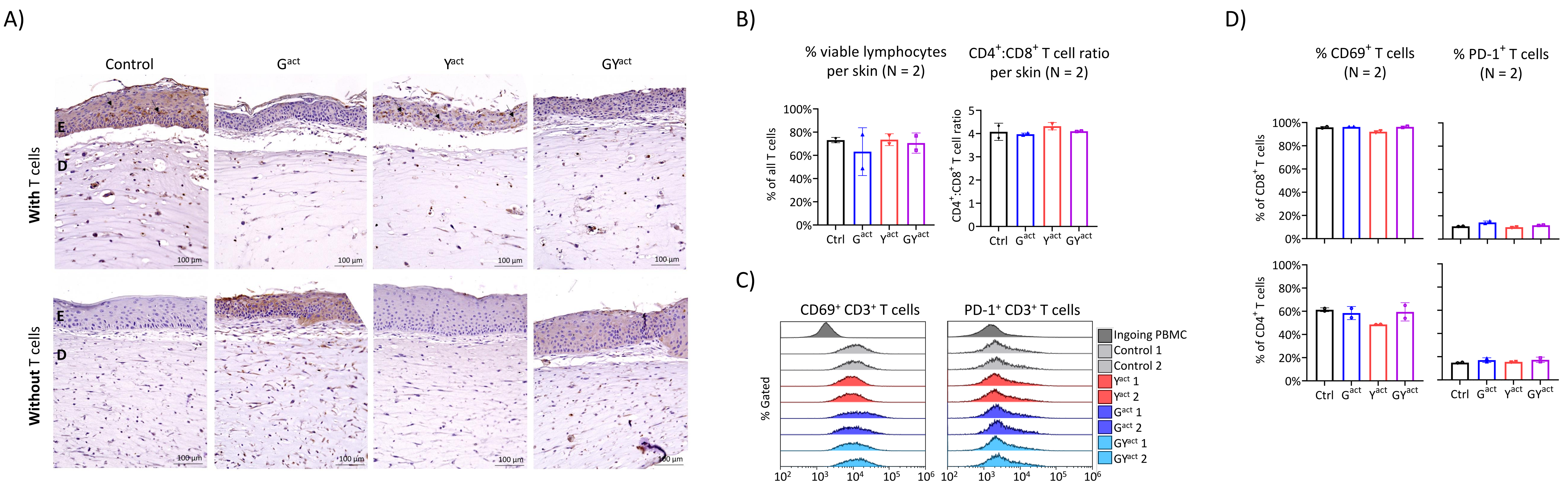

### Stockmaier et al., suppl. Figure S5

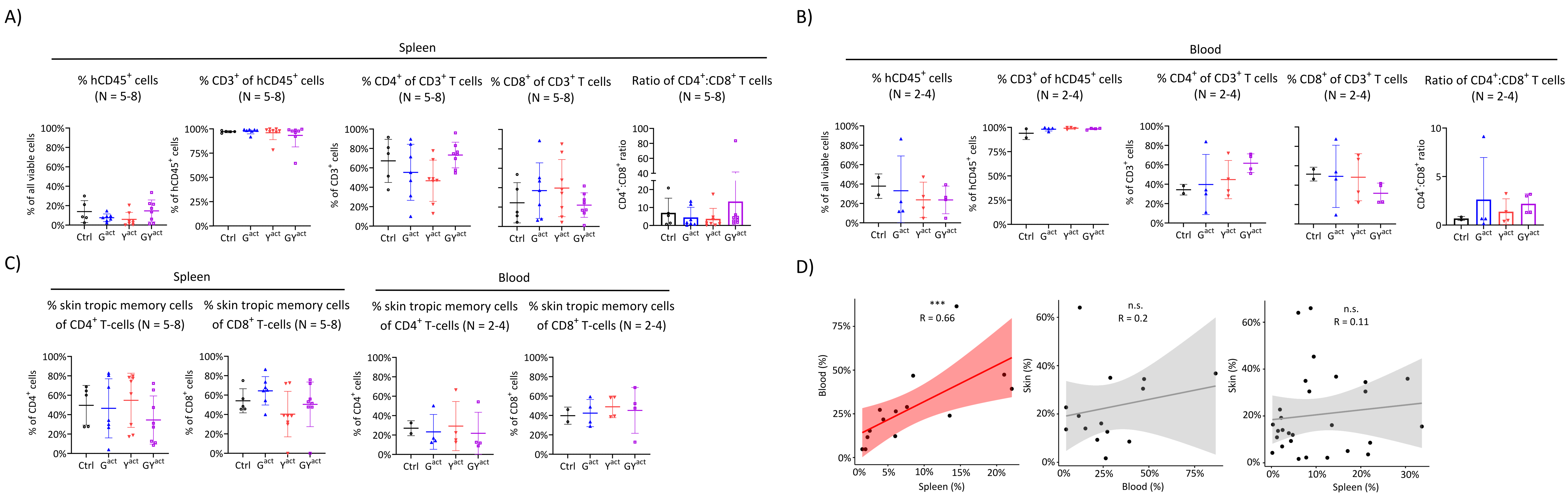
